# Laboratory and Molecular Surveillance of Paediatric Typhoidal *Salmonella* in Nepal: Antimicrobial Resistance and Implications for Vaccine Policy

**DOI:** 10.1101/250142

**Authors:** Carl D Britto, Zoe A Dyson, Sebastian Duchene, Michael J Carter, Meeru Gurung, Dominic F Kelly, David Murdoch, Imran Ansari, Stephen Thorson, Shrijana Shrestha, Neelam Adhikari, Gordon Dougan, Kathryn E Holt, Andrew J Pollard

## Abstract

**Background:** Children are substantially affected by enteric fever in most settings with a high burden of the disease, which could be due to immune naivety, or enhanced risk of exposure to the pathogen. Although Nepal is a high burden setting for enteric fever, the bacterial population structure and transmission dynamics are poorly delineated in young children, the proposed target group for immunization programs.

**Methods:** Blood culture surveillance amongst children aged 2 months to 15 years of age was conducted at Patan Hospital between 2008 and 2016. A total of 198 *S.* Typhi and 66 *S.* Paratyphi A isolated from children treated in both inpatient and outpatient settings were subjected to whole genome sequencing and antimicrobial susceptibility testing. Demographic and clinical data were also collected from the inpatients. The resulting data were used to place these paediatric Nepali isolates into a worldwide context, based on their phylogeny and carriage of molecular determinants of antimicrobial resistance (AMR).

**Results:** Children aged ≤4 years made up >40% of the inpatient population. The majority of isolates (78 %) were *S*. Typhi, comprising several distinct genotypes but dominated by 4.3.1 (H58). Several distinct *S.* Typhi genotypes were identified, but the globally disseminated *S*. Typhi clade 4.3.1 (H58) dominated. The majority of isolates (86%) were insusceptible to fluoroquinolones. This was mainly associated with *S*. Typhi H58 Lineage II and *S*. Paratyphi A; non-susceptible strains from these two genotypes accounted for 50% and 25% of all enteric fever cases. Multi-drug resistance (MDR) was rare (3.5% of *S*. Typhi, 0 *S*. Paratyphi A) and restricted to chromosomal insertions of AMR genes in H58 lineage I strains. Comparison to global data sets showed the local *S*. Typhi and *S*. Paratyphi A strains had close genetic relatives in other South Asian countries, indicating regional strain circulation.

**Conclusions:** These data indicate that enteric fever in Nepal continues to be a major public health issue with ongoing inter- and intra-country transmission, and highlights the need for regional coordination of intervention strategies. The absence of a *S.* Paratyphi A vaccine is cause for concern, given its prevalence as an enteric fever agent in this setting, and the large proportion of isolates displaying fluoroquinolone resistance. This study also highlights an urgent need for routine laboratory and molecular surveillance to monitor the epidemiology of enteric fever and evolution of antimicrobial resistance within the bacterial population as a means to facilitate public health interventions in prevention and control of this febrile illness.

## Introduction

As in most developing countries, invasive bacterial infections account for a significant proportion of paediatric morbidity and mortality in Nepal^1,2^. Enteric fever, caused by *Salmonella enterica* serovars Typhi (*S.* Typhi) and Paratyphi A (*S.* Paratyphi A), is the most common cause of bloodstream infection in Nepal^1,2^ and on a global scale causes an estimated 26 million cases of enteric fever annually of which a large proportion are in children^3,4^. In Nepal, it is estimated that 13% of febrile paediatric cases attending outpatient care are blood culture positive for *S.* Typhi or Paratyphi A^2^. Single nucleotide polymorphism (SNP) genotyping of *S.* Typhi isolated in a study of paediatric enteric fever cases at Patan Hospital in Kathmandu, Nepal during 2005 and 2006 suggested that, among children treated as inpatients, those aged ≤4 years were susceptible to a wider range of haplotypes due to immune naivety^5^. The most common genotype was H58 lineage II (70%), followed by H42 (19%)^5^. Another study of adults and children at Patan Hospital from 2005 to 2009 found that 26% of culture-positive cases were associated with *S*. Paratyphi A; the rest were caused by *S*. Typhi, mainly H58 lineage II (61%) or other H58 (3%), or H42 (15%)^6^. More recently, whole genome sequencing (WGS) was applied to study *S.* Typhi isolates collected during a randomized controlled trial of gatifloxacin vs ceftriaxone for treatment of blood culture confirmed enteric fever at Patan Hospital between 2011 and 2014, and found the H58 genotype continued to dominate the circulating *S.* Typhi population (83%)^7^.

Multi-drug resistant (MDR) *S.* Typhi, defined as resistant to the first-line antibiotics ampicillin, chloramphenicol and co-trimoxazole, became common in the Indian subcontinent in the 1990s^8^, driven by the spread of H58 carrying an IncHI1 plasmid harbouring a suite of antimicrobial resistance (AMR) genes^9^. These *S.* Typhi strains are still circulating in the region, including in India, Pakistan and Bangladesh. In this setting there is also evidence of migration of the AMR genes to the *S*. Typhi chromosome, and acquisition of additional resistance to fluoroquinolones and third-generation cephalosporins, which further limits treatment options in the region^10^. However in Nepal, the MDR H58 *S.* Typhi appears to have been replaced by non-MDR H58 *S.* Typhi carrying the S83F mutation in *gyrA* and other mutations in the quinolone resistance determining region (QRDR) associated with reduced susceptibility to fluoroquinolones^5,6^; and more recently the introduction of fluoroquinolone resistant H58 *S.* Typhi, likely from India, resulting in failure of gatifloxacin treatment^7^. *S*. Paratyphi A in Nepal is generally not MDR, but frequently carries fluoroquinolone non-susceptibility alleles in *gyrA* and *parC.*^11–13^

Given the current treatment complexities of paediatric enteric fever, vaccination would seem the most feasible short-term strategy. There is no vaccine against *S*. Paratyphi A, which accounts for approximately a quarter of disease cases in Nepal. The Vi polysaccharide vaccine against *S*. Typhi is not effective in children under two years of age^14^, and has therefore not been deployed as part of the national immunization schedule in Nepal and is only available privately. While the Vi conjugate vaccines have the potential to reduce the incidence of enteric fever in Nepal, the immunization approach and schedule needs to be clearly defined. This study sheds light on the age distribution of affected inpatient children at Patan Hospital, and the molecular structure and AMR determinants of circulating bacterial pathogen populations causing paediatric enteric fever from 2008 to 2016 in Nepal, with the view of informing preventive strategies including vaccine policy.

## Methods

### Ethics statement

Ethical approval was obtained from the Oxford Tropical Research Ethics Committee (OxTREC) as well as local institutional approval from the Nepal Health Research Council (R31579/CN007).

### Study Setting

Nepal is a low income^15^, landlocked Himalayan nation with an under-five year old mortality rate of 35.8 per 1000 live births as of 2015^16^. Kathmandu Valley, the main urban centre of Nepal, has three districts and a population of 2.5 million^17^ (average population density: 2,372/km^2^) of which 31% are between 0-14 years under age^18^. Over the course of the study, the Patan Academy of Health Sciences (PAHS) was one of only two large hospitals in Kathmandu Valley with referral and paediatric intensive care services. Patan Hospital accepts patients from all over the Valley. Annually the paediatric department cares for over 50,000 outpatients (21% of all hospital outpatient attendances) and accepts approximately 2,700 inpatient admissions. Only 10% of the patients reside outside Kathmandu Valley.

### Surveillance of culture confirmed enteric fever amongst inpatients

Febrile children under 14 years of age, attending PAHS with clinical suspicion of invasive bacterial disease between January 2008 and December 2016 were included in an invasive bacterial disease database as described previously^19^. Inclusion criteria were: clinical presentation indicating an invasive bacterial infection requiring inpatient care with intravenous antibiotics. Blood culture was conducted as described below. Of the patients included in the database, all those that had blood cultures positive for *S*. Typhi or *S*. Paratyphi A were included in the present study, along with relevant demographic data. A random collection of 67 *S*. Typhi isolates and 17 *S*. Paratyphi A isolates were selected for whole genome sequencing; these represent isolates associated with the severe spectrum of paediatric enteric fever presenting to the hospital.

### Isolates collected from outpatients

Children with milder clinical presentations who are usually treated with oral antibiotics as outpatients were not included in the invasive bacterial disease database; however they are subjected to the same microbiological diagnostic procedures as inpatients (as detailed below). A total of 1283 *S*. Typhi and 926 *S*. Paratyphi A isolates from paediatric outpatients were stored between 2008 and 2016; every 10^th^ *S*. Typhi isolate and every 5^th^ *S*. Paratyphi A isolate were included in this current study, representing isolates associated with milder presentation of paediatric enteric fever at the hospital.

### Blood culture processing

Aerobic blood culture bottles were used to culture 3-5 mL of blood, which were then incubated in a BD Bactec FX 40 incubator at 37**°**C for a maximum of 5 days. Turbid samples were then inoculated directly onto MacConkey agar and incubated for maximum of 5 days at 37**°**C to identify potential *S.* Typhi and *S*. Paratyphi A colonies. Candidate *S.* Typhi and *S.* Paratyphi A isolates were further subjected to standard biochemical tests for additional confirmation^20^.

### Antimicrobial susceptibility testing

Antimicrobial susceptibility profiles were gauged by Kirby-Bauer disk diffusion tests. The CLSI (Clinical and Laboratory Standards Institute) guidelines were used to evaluate zones of inhibition for chloramphenicol, co-amoxiclav, co-trimoxazole, cefexime, ceftriaxone, azithromycin, nalidixic acid, and ciprofloxacin^21^. Isolates displaying sensitivity to the tested antimicrobials as per the cut-off values in the CLSI guidelines were designated as susceptible and those that were intermediate (I) or resistant (R) to the tested antimicrobials were designated as insusceptible.

### Genome sequencing and SNP analysis

Briefly, DNA was extracted using the Wizard Genomic DNA Extraction Kit (Promega, Wisconsin, USA), according to manufacturers instructions. Genomic DNA was then subjected to indexed whole genome sequencing on an Illumina Hiseq 2500 platform at the Wellcome Trust Sanger Institute to generate paired-end reads of 100-150 bp in length.

For analysis of SNPs in *S*. Typhi, Illumina reads were mapped to the reference genome sequence of strain CT18^22^ (accession AL515582) using the RedDog (V1beta.10.3) mapping pipeline, available at https://github.com/katholt/RedDog.

RedDog uses Bowtie (v2.2.9)^23^ to map reads to the reference sequence; uses SAMtools (v1.3.1)^24^ to identify SNPs with phred quality scores above 30; filters out those supported by <5 reads or with >2.5 times the average read depth (representing putative repeated sequences), or with ambiguous consensus base calls. For each SNP that passed these criteria in any one isolate, consensus base calls for the SNP locus were extracted from all genomes (ambiguous base calls and those with phred quality scores less than 20 were treated as unknowns and represented with a gap character). These SNPs were used to assign isolates to previously defined lineages according to an extended *S.* Typhi genotyping framework^25^ (code available at https://github.com/katholt/genotyphi). For phylogenetic analyses, SNPs with confident homozygous allele calls (i.e. phred score >20) in >95% of the *S.* Typhi genomes (representing a ‘soft’ core genome of common *S.* Typhi sequences) were concatenated to produce an alignment of alleles at 233,527 variant sites. SNPs called in phage regions, repetitive sequences (354 kb; ∼7.4% of bases in the CT18 reference chromosome, as defined previously) or in recombinant regions identified using Gubbins (v2.0.0)^26^ were excluded, resulting in a final set of 2,187 SNPs identified in an alignment length of 4,809,037 bp for the 198 novel Nepali *S*. Typhi isolates. SNP alleles from *S*. Paratyphi A strain AKU_12601^27^ (accession FM200053) were also included as an outgroup to root the tree.

To provide regional context, genome data from: (i) a published study of mainly Nepali adults^7^ (n=95), (ii) a global *S*. Typhi genome collection^25^ (n=1,221); were subjected to SNP calling and genotyping, resulting in an alignment of 12,216 SNPs for a total of 1,514 isolates. Details and accession numbers of sequence data included in our analysis have been included in **Supplementary Tables 1 & 2**. An additional analysis of all 261 H58 (genotype 4.3.1) from Nepal was carried out in the same manner, resulting in an alignment of 631 SNPs.

To characterize and analyse the genomes of the 66 *S.* Paratyphi A strains, a similar bioinformatic process was adopted using *S.* Paratyphi A AKU_12601^27^ (accession no: FM200053) as the reference genome to create an alignment with another selected 176 isolates from previous studies^28–30^, for global context resulting in an alignment of 5,277 SNPs in a total of 242 *S.* Paratyphi isolates, with alleles from *S.* Typhi CT18^22^ (accession no: AL515582) included as an outgroup to root the tree.

### Phylogenetic analysis

Maximum likelihood (ML) phylogenetic trees were inferred from SNP alignments using RAxML (v8.1.23)^31^, with the generalized time-reversible model, a Gamma distribution to model site-specific rate variation (the GTR+ Γ substitution model; GTRGAMMA in RAxML), and 100 bootstrap pseudo-replicates to assess branch support. The resulting trees were visualized using Microreact^32^ and the R package ggtree^33^. For visualization purposes, *S*. Typhi isolates representing ‘outbreaks’ (defined as members of the same monophyletic clade, isolated from the same study location in the same year) were manually thinned to a single representative.

### Temporal analysis

To investigate the temporal signal and emergence dates of antimicrobial resistance determinants for Nepali *S*. Typhi 4.3.1, we used several methods. First, we used TempEst (v1.5)^34^ to assess temporal structure (i.e. whether the data have clocklike behavior) by conducting a regression of the root-to-tip branch distances of the Nepal H58/4.3.1 ML tree as a function of the sampling time, using the heuristic residual mean squared method with the best-fitting root selected. The resultant data were then visualized in R^35^. To estimate divergence times we analysed the sequence data in BEAST2 (v2.4.7)^36^. We used both constant-coalescent population size and Bayesian skyline tree priors, in combination with either a strict molecular clock model or a relaxed (uncorrelated lognormal distribution) clock model to identify the model that best fits the data. For the BEAST2 analysis the GTR+ Γ substitution model was selected, and the sampling times (tip dates) were defined as the year of isolation to calibrate the molecular clock. For all model and tree prior combinations, a chain length of 100,000,000 steps sampling every 5000 steps was used^37^. The relaxed (uncorrelated lognormal) clock model, which allows evolutionary rates to vary among branches of the tree together with the skyline demographic model, proved to be the best fit for the data. To assess the signal of these Bayesian estimates we conducted a date-randomization test whereby sampling times were assigned randomly to the sequences, and the analysis re-run 20 times^37,38^. These randomization tests were conducted with the same ‘best fit’ models (uncorrelated lognormal clock and skyline demographic). This test suggested that the data display ‘strong’ temporal structure^37^.

For the final analysis reported here, 5 independent runs conducted with a chain length of 600,000,000 states, sampling every 300,000 iterations, were combined using LogCombiner (v2.4.7)^36^ following removal of the first 10% of steps from each as burn-in. Maximum-clade credibility (MCC) trees were generated with ‘keep target heights’ specified for node heights using TreeAnnotator (v2.4.7)^36,39^. The effective sample sizes from the combined runs were estimated to be >200 for all reported parameters.

### In silico resistance plasmid and AMR gene analysis

The mapping based allele typer SRST2^40^ was used to detect plasmid replicons and acquired AMR genes and determine their precise alleles, by comparison to the ARG-Annot^41^ and ResFinder^42^ databases (for AMR genes) and PlasmidFinder^41^ (for plasmid replicons). Where AMR genes were observed without evidence of a known resistance plasmid, raw read data was assembled *de novo* with SPAdes (v3.7.1)^43^ and Unicycler (v0.3.0b)^44^ and examined visually using the assembly graph viewer Bandage (0.8.1)^45^ to inspect the composition and insertion sites of resistance-associated transposons. These putative transposon sequences were annotated using Prokka (v1.11)^45^ followed by manual curation, and visualized using the R package *genoPlotR*^45^. SNPs in the QRDR of *gyrA, gyrB, parC* and *parE* genes, which are associated with reduced susceptibility to fluoroquinolones in *S.* Typhi, *S*. Paratyphi A and other species^7^, were extracted from the whole genome SNP alignments.

### Nucleotide sequence and read data accession numbers

Raw sequence data have been deposited in the European Nucleotide Archive under project PRJEB14050; and individual accession numbers are listed in **Supplementary Tables 1–2.** Genome assemblies for isolates RN2293 and RN2370 were deposited in GenBank.

## Results

### Paediatric enteric fever surveillance

Blood cultures were performed on 11,430 children with a suspected invasive bacterial infection and requiring inpatient care with intravenous antibiotics and supportive care. Of these, 129 had blood cultures positive for the enteric fever agents *S*. Typhi (n=102, 79%) or *S*. Paratyphi A (n=27, 21%). Relevant patient characteristics are reported in **Table 1**. Most cases of culture-confirmed enteric fever (n=83, 64%) occurred between the hot and rainy months of May and October. However, a substantial proportion (36%) of cases also occurred in colder months, indicating perennial transmission. Children under 5 years of age accounted for 45% of the disease burden among inpatients, with children under 2 years of age accounting for 18% (**Table 1**). Clinical suspicion of enteric fever at presentation was significantly lower amongst children under 2 years with culture-confirmed infection (13% vs. 52%, p=0.0005 using Fisher’s exact test; **Table 1**), highlighting the undifferentiated febrile nature of the disease even in an endemic setting such as Nepal.

**Table 1.** Hospital based (inpatient) paediatric enteric fever surveillance

|  |  |  |  |  |
| --- | --- | --- | --- | --- |
| Total blood cultures performed | 11430 |  |  |  |
| Total number of significant cultures | 1048 (9.2%) |  |  |  |
| Total number of enteric fever pathogens | 129 (1.1%) |  |  |  |
| *S. Typhi* | 102 (0.9%) |  |  |  |
| *S. Paratyphi A* | 27 (0.2%) |  |  |  |
| Age stratified characteristics of blood-culture positive enteric fever patients |  |  |  |  |
| Age groups | <2 y | 2-4 y | 5-9 y | 10-14 y |
| Number | 23 | 35 | 39 | 31 |
| Median age (years) | 1.2 | 3.3 | 6.8 | 11.8 |
| Male (%) | 16 (70%) | 23 (66%) | 24 (62%) | 18 (58%) |
| Median temperature at admission (C°) (range) | 37.2 (36.7-38.9) | 38.3 (36.5-39.9) | 38.9 (36.1-40.5) | 37.2 (36.5-39.2) |
| Median duration of admission (days) (range) | 6 (2-23) | 6.5 (1-19) | 8 (2-36) | 7.5 (3-20) |
| Enteric fever suspicion on admission (%) | 3 (13%) | 17 (49%) | 19 (49%) | 19 (61%) |

### Phylogenetic structure of paediatric isolates from Nepal

The genomes of *S*. Typhi isolated from inpatient surveillance (n=67) and a random selection of isolates from outpatients (n=131) were sequenced and subjected to SNP genotyping and phylogenomic analysis as described in **Methods**. The resulting phylogeny (**Figure S1**) revealed the presence of 8 distinct genotypes, each corresponding to a different subclade including 2.0.0 (N=1, 0.5%) 2.2.0 (N=10, 5%), 2.3.4 (N=2, 1%), 3.2.2 (N=6, 3%), 3.3.0 (N=19, 9.6%), 3.3.1 (N=3, 1.5%), 4.1.0 (N=3, 1.5%), and 4.3.1 (N=154, 77.8%). There was no significant association between genotype and treatment status (outpatient vs. inpatient), period of isolation (**Figure 1A**) or patient age (**Figure 1B**).

**Figure 1:**
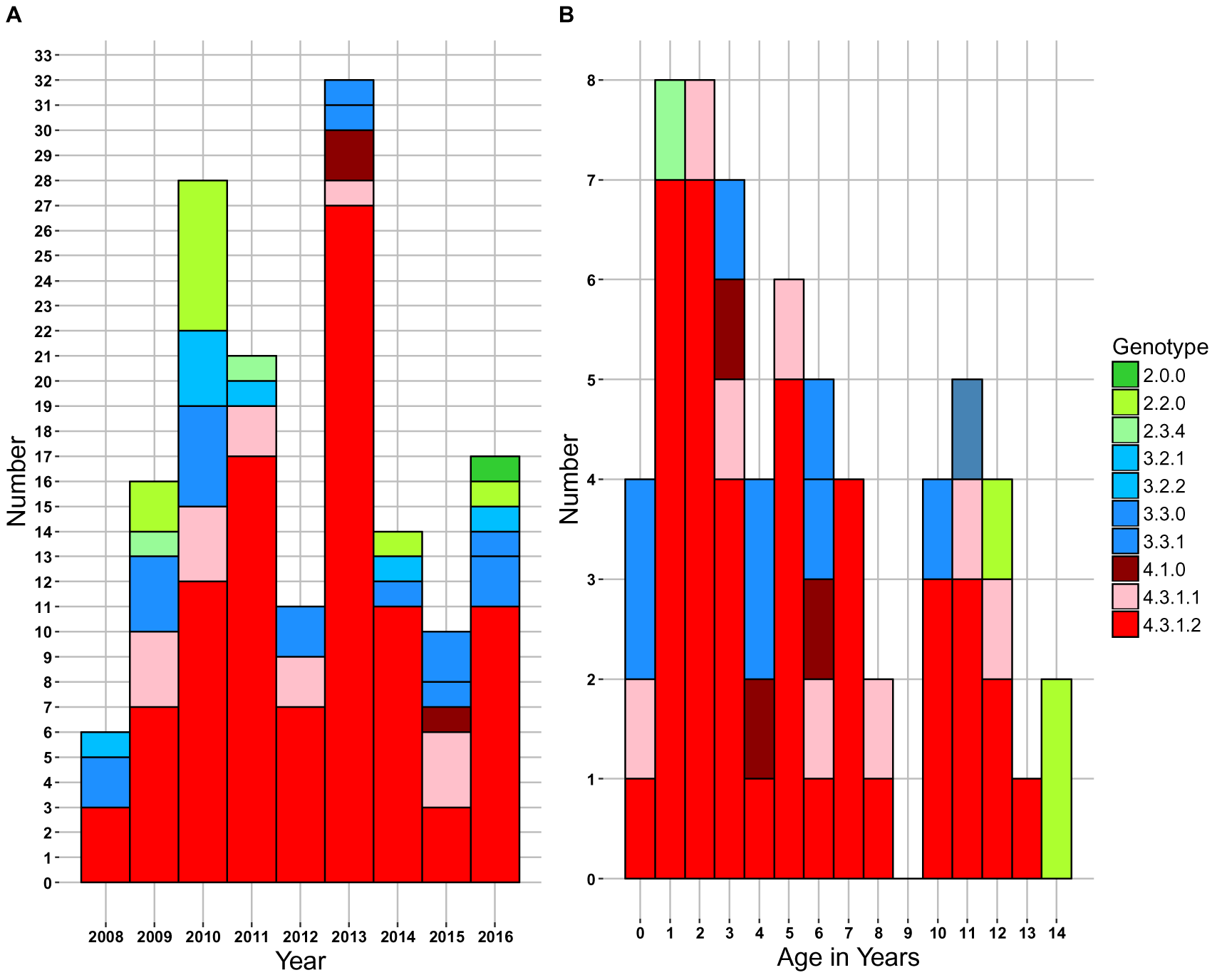
Nepal peadiatric *S.* Typhi genotypes. (A) Genotypes observed per annum. (B) Genotypes observed per age in years. Individual *S.* Typhi genotypes are coloured as described in the inset legend.

To place the novel paediatric isolates in context, we constructed a whole genome phylogeny including other *S.* Typhi previously sequenced from adults in Nepal, and a global collection of *S*. Typhi (**Figure 2**; an interactive version of the phylogeny and associated geographical data are also available for exploration online at https://microreact.org/project/SJmU6dhlz). The novel paediatric isolates clustered together with the adult isolates from Nepal, with no evidence of certain genotypes circulating in children more so than adults. In comparison to global isolates, Nepali isolates clustered with those from other regions in the Indian subcontinent, suggesting ongoing transmission within the region (**Figure 2**); indeed 14% of the novel Nepali paediatric isolates and 15% of the previously sequenced Nepali isolates were closest to an isolate from outside Nepal (majority from neighbouring India, Bangladesh or Pakistan), indicating frequent pathogen transfer within the region.

**Figure 2:**
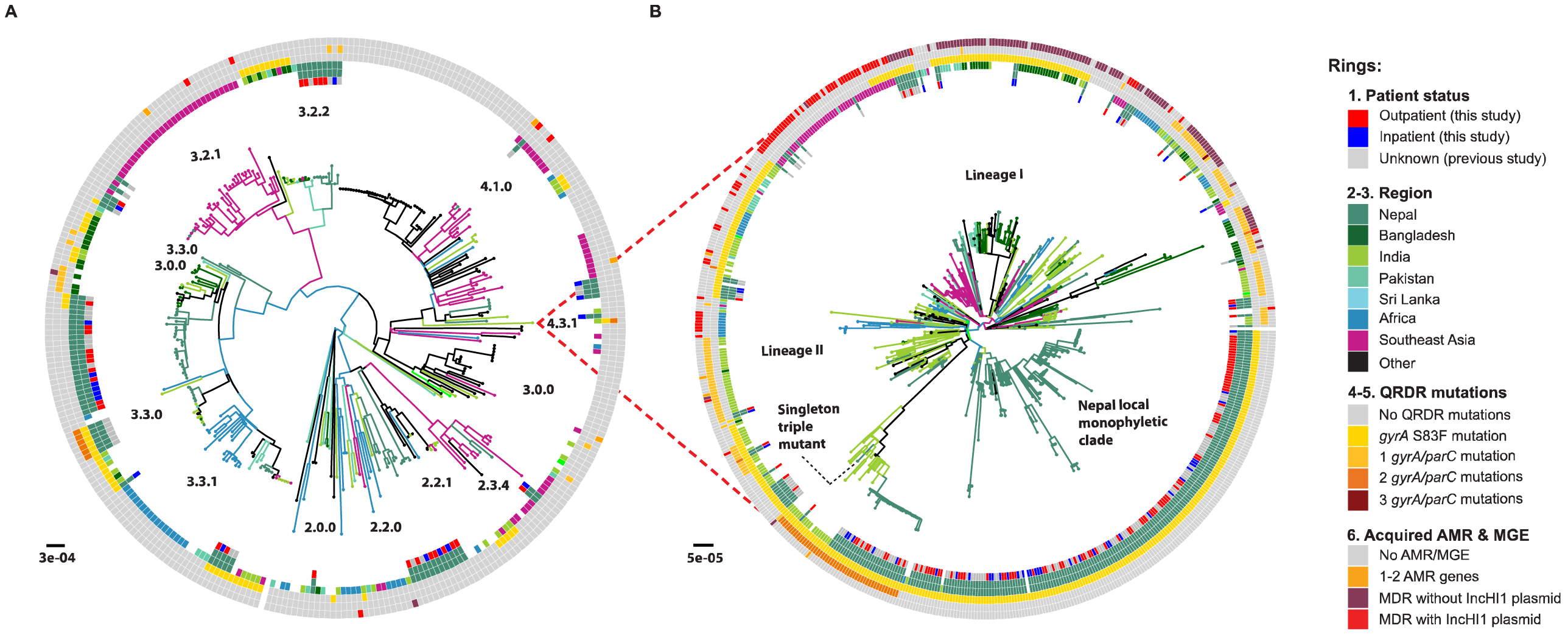
Global population structure of Nepalese *S.* Typhi genotypes. (A) Global population structure of Non-H58 (4.3.1) Nepalese genotypes. (B) Global population structure of H58 (4.3.1). Inner ring shows patient status. Branch colours indicate country/region of origin, as do the second ring and third rings from the inside. Fourth ring from the inside indicates QRDR *gyrA* S83F mutation. Fifth ring from the inside indicates the number of *gyrA* and *parC* QRDR mutations. Outer ring describes the presence of MDR. All rings and branches are coloured as per the inset legend. Branch lengths are indicative of the estimated number of substitutions rate per variable site, and the tree was outgroup rooted with a *S.* Paratyphi A strain

We used the same approach to investigate genome variation amongst 66 *S*. Paratyphi A isolated from inpatients (n=17) and outpatients (n=49) in Nepal, in the context of globally representative genome diversity (**Figure 3**; interactive version available at https://microreact.org/project/rk2ec5mWM). The Nepali *S.* Paratyphi A population was far less diverse than that of *S*. Typhi; most belonged to lineage A and clustered into two distinct subgroups, which we designated sublineages A1 and A2 (see **Figure 3**). Akin to *S*. Typhi, the global context of *S.* Paratyphi A also revealed close clustering with isolates from other regions in the Indian subcontinent and China, which where *S*. Paratyphi A infections occur at high prevalence.

**Figure 3:**
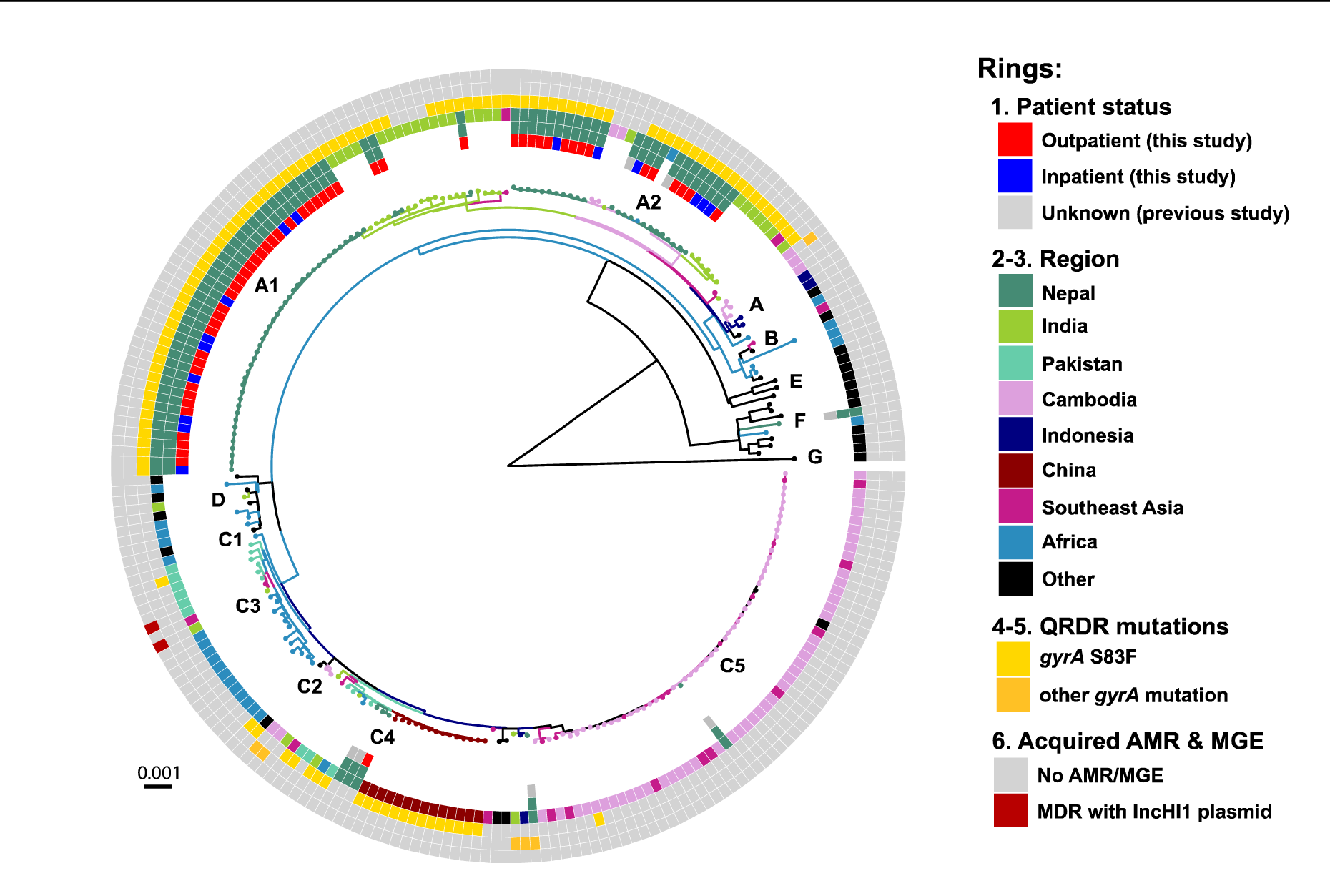
Global population structure of *S*. Paratyphi A. Inner ring indicates patient status. Branch colours indicate country/region of origin, as do the second and third rings from the inside. Fourth ring from the inside indicates QRDR *gyrA* S83F mutation. Fifth ring from the inside indicates the number of additional *gyrA* and *parC* QRDR mutations. Outer ring describes the presence of MDR. All rings and branches are coloured as per the inset legend. Branch lengths are indicative of the estimated number of substitutions rate per variable site, and the tree was outgroup rooted with *S.* Typhi strain CT18

### Antimicrobial resistance (AMR)

Amongst the paediatric isolates analysed in this study, most *S*. Typhi isolates (96%) and all *S.* Paratyphi A were susceptible to traditional first-line antibiotics cotrimoxazole, ampicillin and chloramphenicol (**Figure 4**). Most (86%) of *S*. Typhi and all the *S.* Paratyphi A of isolates were insusceptible to the fluoroquinolone ciprofloxacin (assessed by disk diffusion; **Figure 4**). MDR was observed in six *S*. Typhi (3%) and no *S*. Paratyphi A. There were no differences in the frequency of MDR or fluoroquinolone insusceptibility between the paediatric inpatients and outpatients (OR for MDR = 0.97, 95% CI 0.23 – 4.00; and OR for fluoroquinolone insusceptibility = 1.23, 95% CI 0.62 – 2.44).

**Figure 4:**
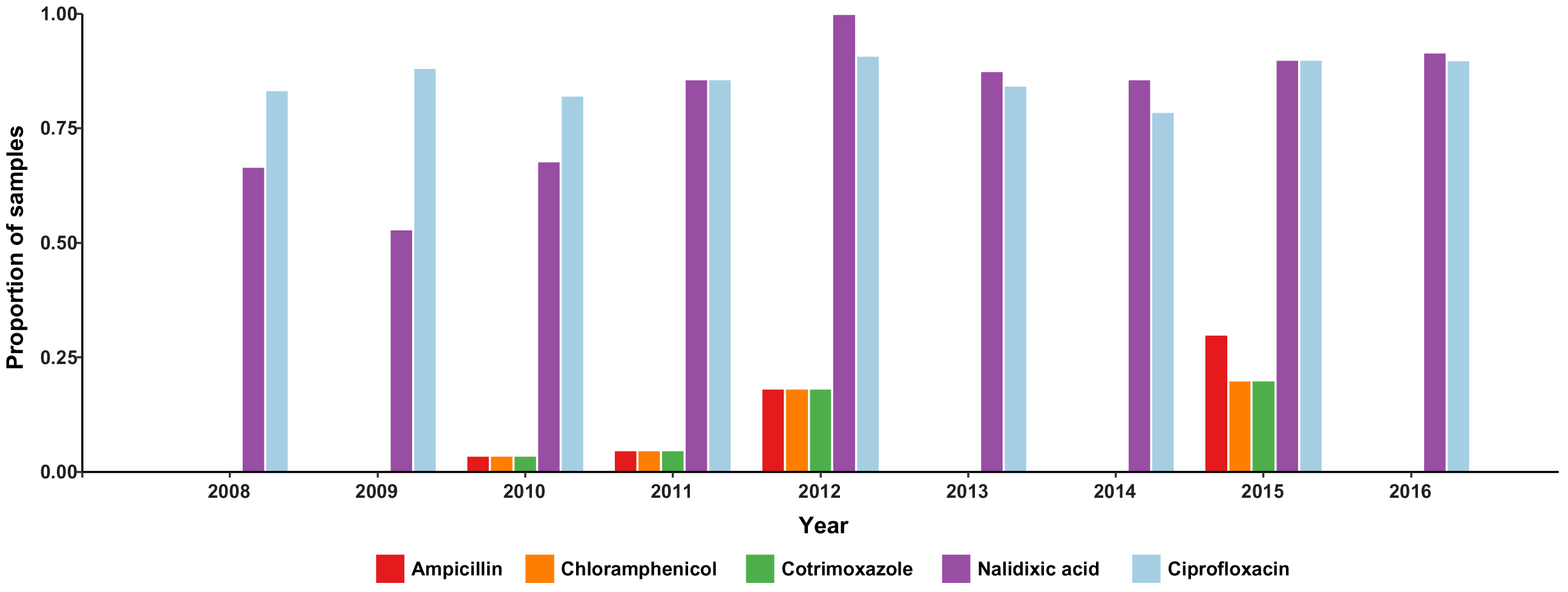
Bars are coloured as described in the inset legend. Susceptibility to Ampicillin, Chloramphenicol, Ciprofloxacin, Cotrimoxazole, Nalidixic acid, Cephalosporin, Ceftriaxone, Cefixime, and Azithromycin were tested. No resistance to Azithromycin, Ceftriaxone, and Cefixime was observed.

Genetic determinants of AMR detected in the paediatric isolates are summarized in **Table 2**. All *S.* Paratyphi A (besides the single lineage C4 isolate) carried the *gyrA* S83F mutation responsible for nalidixic acid resistance and fluoroquinolone insusceptibility. *S*. Typhi isolates displaying fluoroquinolone insusceptibility harboured known QRDR SNPs (**Table 2**); these included isolates of genotypes 4.3.1 (*gyrA* SNPs), 3.3.0 (*parE* SNPs), and 3.3.1 (*gyrA* and *parE* SNPs, see **Figure S1**). Sixteen *S*. Typhi isolates (all genotype 4.3.1) were QRDR ‘triple mutants’, which are associated with failure to respond to fluoroquinolone therapy^7^. All MDR isolates (n=6) belonged to *S*. Typhi genotype 4.3.1 and harboured the acquired AMR genes *catA*, *dfrA7, sul1*, *sul2*, *strA, strB* and *bla_TEM-1_*, conferring resistance to chloramphenicol, co-trimoxazole, streptomycin and ampicillin. An additional genotype 4.3.1 isolate carried a subset of four of these genes (*sul2*, *strA, strAB* and *bla_TEM-1_*) and displayed resistance to ampicillin but was sensitive to co-trimoxazole and chloramphenicol (consistent with the lack of *dfr* and *cat* genes). Acquired AMR genes were not detected amongst the *S*. Paratyphi A.

**Table 2.** Genetic determinants of antimicrobial resistance in paediatric isolates from Nepal

|  | *S. Typhi* | *S. Paratyphi A* |
| --- | --- | --- |
| **Total isolates** | **198** | **66** |
| **QRDR** | **164 (82.8%)** | **65 (98%)** |
| *gyrA* S83F | 143 (72%) | 65 (98%) |
| *gyrA* S83F only | 15 (7.6%) | 0 |
| *gyrA* S83F, *gyrA* D87N | 16 (8.1%) | 0 |
| *gyrA* S83F, *gyrA* D87N, *parC* S80I | 15 (7.6%) | 0 |
| *gyrA* S83F, *gyrA* D87N, *parC* E84G | 1 (0.5%) | 0 |
| *gyrA* S83F, *parC* E84G | 1 (0.5%) | 0 |
| *gyrA* S83F, *parE* A364V | 5 (2.5%) | 0 |
| *gyrA* S83Y only | 6 (3%) | 0 |
| *parE* A364V only | 15 (7.6%) | 0 |
| **Acquired AMR genes** | **7 (3.5%)** | **0** |
| *blaTEM-1*, *strAB*, *sul2* | 1 (0.5%) | 0 |
| *catA1*, *dfrA7*, *sul1*, *blaTEM-1*, *strAB*, *sul2* (+ *gyrA* S83F) | 4 (2%) | 0 |
| *catA1*, *dfrA7*, *sul1*, *blaTEM-1*, *strAB*, *sul2* (+ *gyrA* S83Y) | 2 (1%) | 0 |

The full suite of seven acquired AMR genes are common amongst *S*. Typhi globally and are typically located within a composite transposon, comprising Tn*6029* (*sul2*, *strA, strAB* and *bla*^*TEM-1*^) and Tn*21* (*dfrA7, sul1*) inserted within Tn*9* (*catA*), which is most often carried on IncHI1 plasmids^9^. Here, all MDR isolates carried this typical composite transposon, inserted in the chromosome between genes STY3618 and STY3619 and associated with an 8 bp target site duplication (GGTTTAGA), consistent with integration mediated by the flanking IS*1* transposases of Tn*9* (see **Figure 5**). The additional ampicillin resistant isolate carried only transposon Tn*6029*, which was inserted directly into the chromosomal pseudogene *slrP* and associated with an 8 bp target site duplication (TAGCTGAT), consistent with integration mediated by the flanking IS*26* transposases of Tn*6029*.

**Figure 5:**
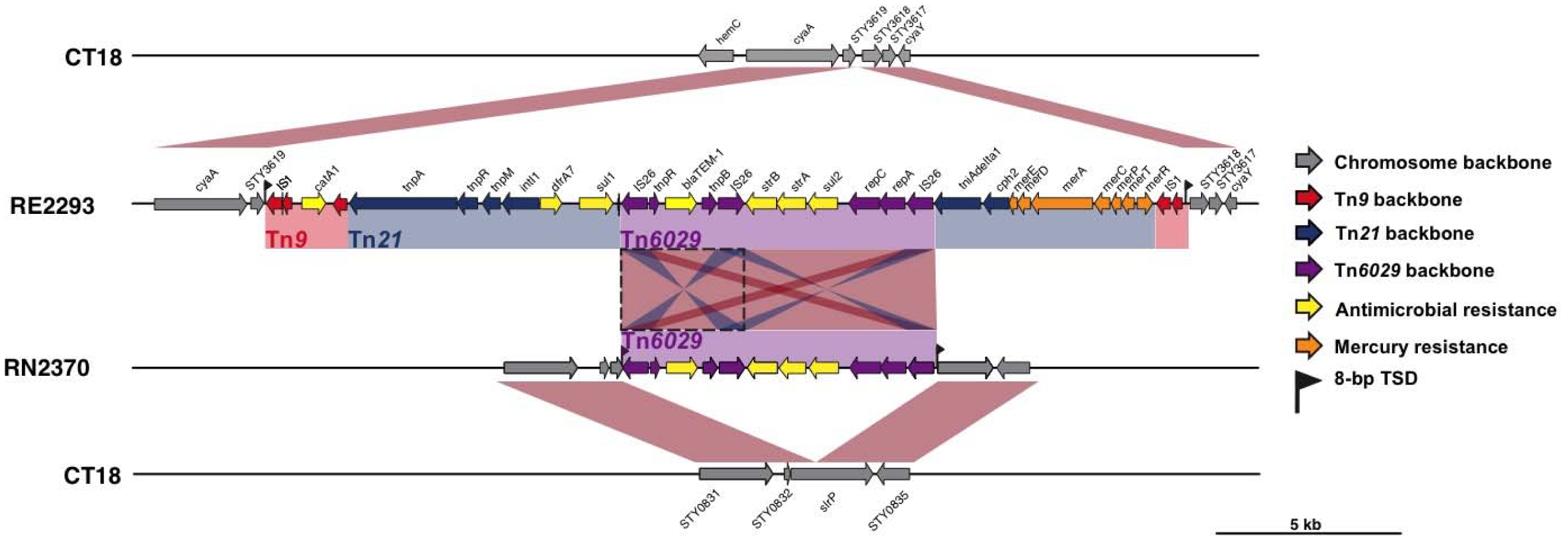
Insertion sites of transposons observed in *S.* Typhi from Nepal. Genes and transposons are indicated according to the inset legend. TSD indicates target site duplication, and *Tn* indicates transposon.

### Evolutionary history of AMR S. Typhi 4.3.1 in Nepal

We constructed a dated phylogeny of all available *S*. Typhi 4.3.1 from Nepal, using BEAST2 **Figure 6,** interactive version available at https://microreact.org/project/rJnfyOGxG). This analysis yielded a local substitution rate of 0.8 SNPs per genome per year (95% highest posterior density (HPD), 0.5 – 1.1) or 1.7x10^-7^ genome-wide substitutions per site per year 95% HPD, 1.1x10^-7^ – 2.4x10^-7^). The data showed strong temporal structure to support these results (see **Methods** and **Figure S2**), which were consistent with previous estimates for global *S*. Typhi 4.3.1^10^. We estimated the most recent common ancestor (mrca) for all *S*. Typhi 4.3.1 in Nepal existed circa 1993, similar to the mrca estimated globally for *S*. Typhi 4.3.1, which is predicted to have emerged in neighbouring India^10^.

**Figure 6:**
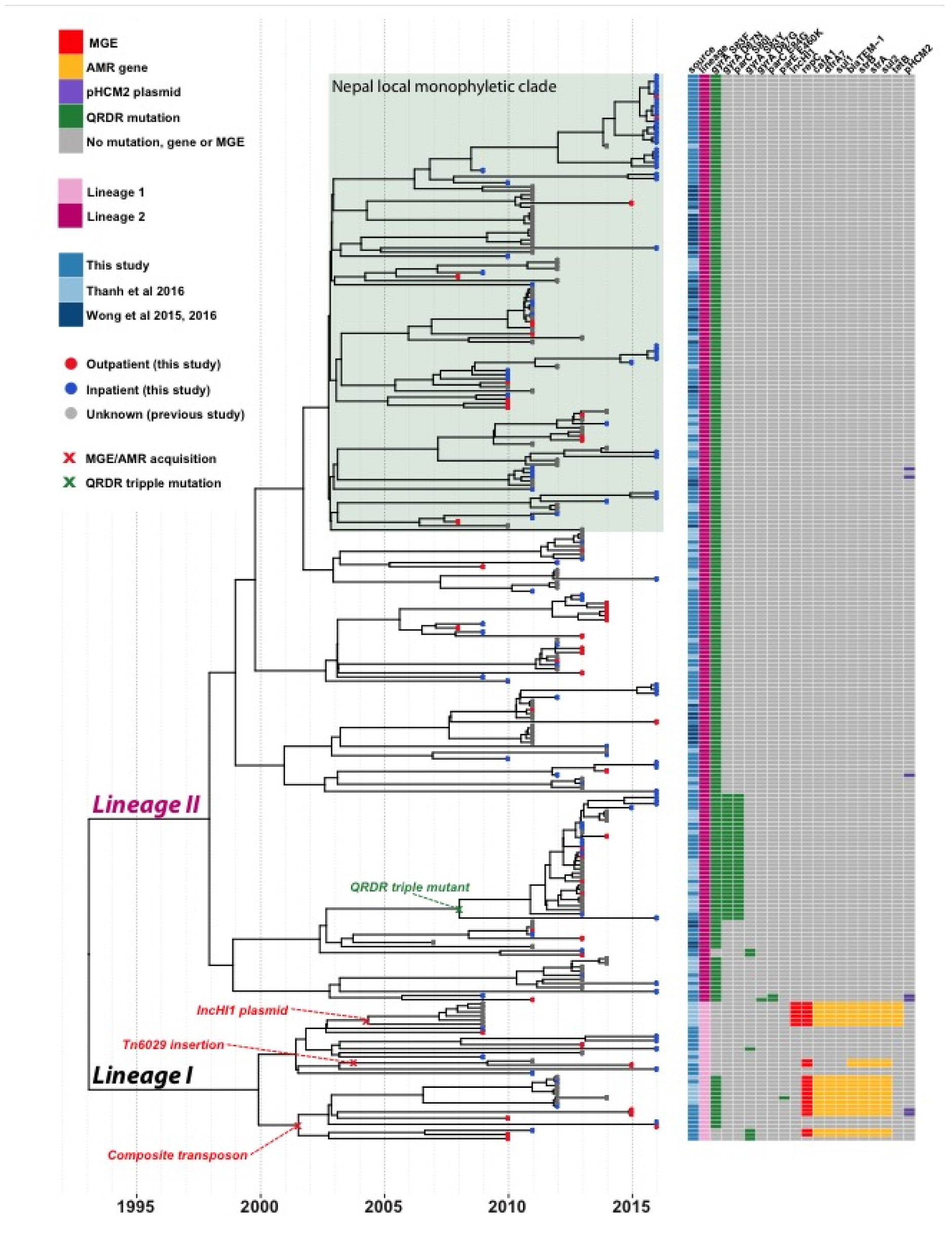
BEAST2 maximum-clade credibility phylogenetic tree of Nepalese genotype 4.3.1 (H58) *S.* Typhi. Tips colours indicate outpatient (red) vs. inpatient (blue) isolates. Acquisitions of molecular determinants of resistance are labelled according to the inset legend. A local Nepal monophyletic clade is shaded in green. Branch lengths are indicative of time and the scale bar corresponds to calendar years

Both of the previously described sublineages of *S*. Typhi 4.3.1 (I and II) were present amongst the Nepali isolates, however (i) lineage II was far more common (67% vs. 10% of paediatric isolates from this study; 68% vs 10% of isolates from other studies); and (ii) the lineages were associated with different AMR patterns (**Figure 6**): lineage I was associated with MDR (59% of lineage I vs 0 lineage II, p<1x10^-15^), while lineage II was associated with QRDR mutations (99% of lineage II vs 50% of lineage I, p<1x10^-15^). The majority of isolates formed a local monophyletic clade that was not detected in other countries in the global collection, indicative of local clonal expansion in Nepal. The relative proportion of local *S*. Typhi infections caused by lineage II increased after 2010 (40% pre-2010 vs 74% from 2011 onwards, p=1x10^-7^), suggesting clonal replacement of the MDR-associated Lineage I with the expansion of the quinolone resistance-associated Lineage II over time.

Most of the ciprofloxacin resistant triple mutant isolates harboured *gyrA* S83F, *gyrA* D87N, and *parC* S80I and formed a monophyletic subclade of lineage II, together with those previously reported as associated with gatifloxacin failure during the treatment trial in 2013-2014^7^. We dated the mrca of this subclade to 2008 (95% HPD, 1998–2011; see **Figure 6**), and comparison to the global tree confirmed it most likely originated in India^7^ and was introduced to Nepal at least twice (see **Figure 2B**). We also identified a distinct ciprofloxacin resistant triple mutant (harbouring *gyrA* S83F, *gyrA* D87G, and *parC* E84G) that was isolated rom a five-year old girl in 2011. This was also *S*. Typhi 4.3.1 lineage II but shared no particularly close relatives in the Nepali or global collections (**Figure 2B**, **Figure 6**).

All isolates with acquired AMR genes belonged to Lineage I: one cluster of IncHI1 plasmid-containing isolates (from a previous study conducted by Thanh et al 2016) with a mean tmrca of 2004 (95% HPD, 1996-2007); two related clusters with the composite transposon inserted in the chromosome after STY3618, with mean tmrca 2001 (95% HPD, 1995-2009); and one cluster with Tn*6029* inserted in the chromosome, with mean tmrca 2003 (95% HPD, 1997-2010) (see **Figure 6**).

## Discussion

These data show that there is a substantial burden of enteric fever amongst children in Nepal **Table 1**), the majority of which (86%) is insusceptible to fluoroquinolones (**Table 2**). Genomic analysis revealed substantial diversity within the local pathogen population (**Figure 1 & S1),** with evidence of transfer of *S*. Typhi and *S*. Paratyphi A between Nepal and neighbouring countries in South Asia (**Figures 2 and 3**), and intermingling of isolates from adults and children consistent with transmission across age groups (**Figure 2**). Data from 2005-2006 suggested that younger children were more susceptible to a wider range of genotypes, a phenomenon attributed to a naïve immune response^5^. A decade later this tendency seems to have shifted towards a more pathogen driven trend as seen in **Figure 1**, which shows the *S*. Typhi 4.3.1 genotype is dominant regardless of the age of the host. This is onsistent with recent mathematical modeling of historical enteric fever patterns in this setting, which identified the introduction of AMR 4.3.1, as well as an increase in migration of immunologically naïve 15-25 year olds from outside the Kathmandu Valley, as key drivers of the local typhoid problem^46^.

The high frequency of fluoroquinolone insusceptibility is attributable to indiscriminate and uncontrolled use of antimicrobials, which since the turn of the century have been used to treat range of infections common in the tropics in addition to enteric fever. Fluoroquionolone insusceptibility has been observed locally^7^, associated with mutations in *gyrA* and *parC*. Our data show that the problem of fluoroquinolone insusceptible enteric fever in Nepali children is mainly driven by two locally established pathogen variants, namely *S*. Typhi 4.3.1 (H58) lineage II harbouring the *gyrA*-S83F mutation (accounting for 50% of all enteric fever, 57% of non-susceptible cases, and 66% of all *S*. Typhi) and *S*. Paratyphi A clade A harbouring the *gyrA*-S83F mutation (accounting for 25% of enteric fever, 28% of non-susceptible cases, and 98% of all *S*. Paratyphi A). These strains have been present since the increase in local case numbers began in 1997, and their arrival likely contributed to the increased disease burden^46^. The universal fluroquinolone resistance demonstrated by the *S*. Paratyphi A population is of great concern particularly since a vaccine against paratyphoid fever is still in development.

Notably, the fully fluoroquinolone resistant triple mutant *S*. Typhi strain that was first detected in local adults in 2013 and halted the gatifloxacin treatment trial was still causing in disease in Nepali children in 2015-2016, but was rare (2.5% of cases in 2015-16) and showed no signs of displacing the wider population that carries only the *gyrA*-S83F mutation (65% of cases in 2015-16). This lack of clonal replacement is consistent with the presence of a single, distinct, triple mutant *S*. Typhi strain isolated from a 5-year old girl in 2011, which had no descendant strains detected amongst the 126 cases examined from 2012-16, suggesting it has not spread within the local human population. The lack of fully resistant *S*. Paratyphi A is also notable. It has been shown that the *gyrA*-S83F mutation is not associated with a fitness cost in *S*. Typhi and can be maintained in the absence of direct selection from fluoroquinolones; however our data suggest the same is not true of the triple mutants, hence limiting exposure to fluoroquinolones may at least control the spread of highly resistant strains.

Acquired resistance to other antimicrobials was rare, and in the paediatric population was associated only with *S*. Typhi 4.3.1 lineage I strains carrying chromosomally integrated AMR genes (**Figure 6**). This has not been reported previously in the local population, where MDR *S.* Typhi has typically been associated with plasmids^47^. Here we identified at least two distinct AMR gene integration events, that we estimate occurred contemporaneously with the MDR plasmid circulating in the early 2000s (**Figure 6**). Although similar findings have also been reported from *S.* Typhi strains in other neighbouring countries of India and Bangladesh^10^, this is the first description in strains from Nepal. Notably, in addition to the integration of the typical *S*. Typhi MDR composite transposon mediated by IS*1* transposase genes of Tn*9*, we identified for the first time direct integration of Tn*6029* into the *S*. Typhi chromosome (**Figure 5**), mediated by IS*26* and conferring ampicillin resistance in the absence of resistance to chloramphenicol or co-trimoxazole.

The findings of this study supplement our understanding of enteric fever in an endemic setting. The occurrence of disease in the <5 years population is in agreement with the other multi-centre data from South Asia, underscoring the importance of understanding the disease transmission dynamics and preventive strategies in the vulnerable population. The magnitude of disease occurrence in this age group is still an underestimation for several reasons; clinical suspicion of enteric fever in this age group is generally low as evidenced in these data and this trend has also been reported in other endemic regions^48^. The lack of clinical suspicion leads to a lack of diagnostic testing, which is in itself, fraught with impediments to reliable results. Blood culture, which is the feasible gold standard diagnostic performs poorly in this population owing to the difficulty in obtaining the required amount of blood and due to pre-treatment with antimicrobials prior to obtaining a blood sample. Despite the unique challenges associated with diagnosing enteric fever in this population and the supposed lack of exposure, reports from various endemic regions continue to reiterate the enormous burden of enteric fever in pre-school children. Coupled with the problem of antimicrobial non-susceptibility once a diagnosis is made, these difficulties highlight the urgent need for enteric fever vaccines in children under 5. However vaccination options for these children are limited due to the poor immunogenicity of the Vi polysaccharide vaccine in infants and the difficulty in administering the Ty21a vaccine. Until the Vi conjugate vaccines are rolled out, in addition to improving sanitation and providing clean water, antimicrobial treatment remains the only short-term option for containing the disease in this age group.

Cephalosporins are currently the first-line treatment for enteric fever in Nepal. We did not detect any cephalosporin non-susceptibility in these isolates, however it is anticipated that this will emerge via the acquisition of plasmid-encoded extended-spectrum beta-lactamase genes, as has recently been observed among *S*. Typhi isolates from neighbouring India and Pakistan^49–52^. Given the re-emergence of antimicrobial sensitivity to chloramphenicol and co-trimoxazole as evidenced in this study, it may be logical to shift to these first-line drugs for treating enteric fever; indeed there has already been a case report demonstrating efficacy of co-trimoxazole treatment in the treatment of fluoroquinolone resistant H58 *S*. Typhi in this setting^53^. We acknowledge the possibility that typhoidal *Salmonella* strains will acquire resistance to these antibiotics when re-introduced and the cycling of antimicrobials is seldom sufficient to effectively prevent MDR in the long-term. However we propose this short-term strategy might be commissioned until the typhoid conjugate vaccines are deployed, in order to conserve cephalosporins and macrolides for the treatment of other tropical infections which require higher-end antibiotics.

## Conclusion

These data highlight the burden of enteric fever in children in Nepal while demonstrating the importance of laboratory and molecular surveillance in endemic regions. Those under the age of 5 years contributed most to the burden of enteric fever among inpatients who represent the severe spectrum of disease. The substantial contribution of those less than 2 years emphasize the urgent need for the Vi conjugate vaccine in regions such as Nepal where antimicrobial therapy is currently the main modality against enteric fever. Antimicrobial non-susceptibility continues to complicate management protocols and calls for prudent strategies aimed at conserving the currently effective drugs while buying time for vaccine deployment. Finally, the control of enteric fever in Nepal and South Asia requires a coordinated strategy given the inter-country transmission that occurs with the Indian subcontinent. The Vi conjugate vaccines offer the real possibility of controlling enteric fever but eradication will only become a possibility when the immunization strategy is supplemented by the provision of clean water and improved sanitation.

## Acknowledgements, Funding and Conflicts of Interest

CB is a Rhodes scholar, class of 2015 funded by the Rhodes trust. ZAD was supported by the Wellcome Trust of Great Britain (106158/Z/14/Z). SD was supported by a McKenzie fellowship from the University of Melbourne. KEH was supported by the NHMRC of Australia (Fellowship #1061409). GD is supported by the NIHR Cambridge, BRC and the Wellcome trust. AJP is funded the NIHR Oxford, BRC and Wellcome trust. The authors also wish to acknowledge the Gates foundation, which support enteric fever studies conducted by our group in Nepal. The WGS sequencing of isolates in this study was funded by the Wellcome Trust Strategic Award: “A strategic vision to drive the control of enteric fever through vaccination”.

AJP has previously conducted studies on behalf of Oxford University funded by vaccine manufacturers, but currently does not undertake industry funded clinical trials. AJP chairs the UK Department of Health’s (DH) Joint Committee on Vaccination and Immunisation (JCVI) and is a member of the World Health Organisation Strategic Group of Experts (SAGE); the views expressed in this manuscript do not necessarily reflect the views of JCVI, DH or SAGE. The other authors have no conflicts of interest.

---

**Supplementary Figure 1.**
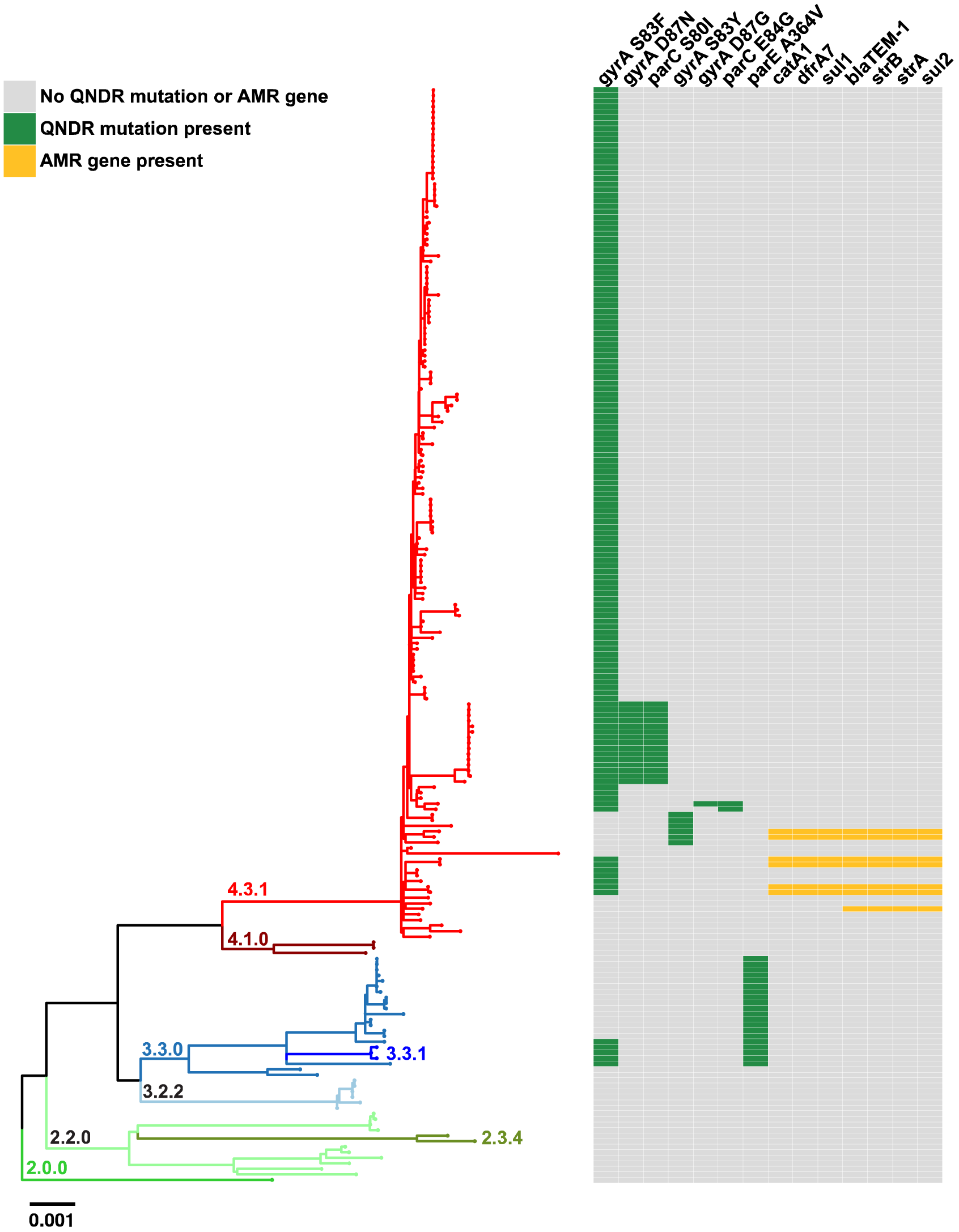
Population structure of paediatric *S*. Typhi in Nepal. Paratyphi A outgroup rooted maximum likelihood phylogeny. Branch colours indicate the genotype (as labelled). Branch lengths are indicative of the estimated number of substitution rate per variable site. The adjacent heatmap shows the presence of AMR genes (yellow) and QRDR mutations (green) present in each isolate as described in the inset legend.

**Supplementary Figure 2.**
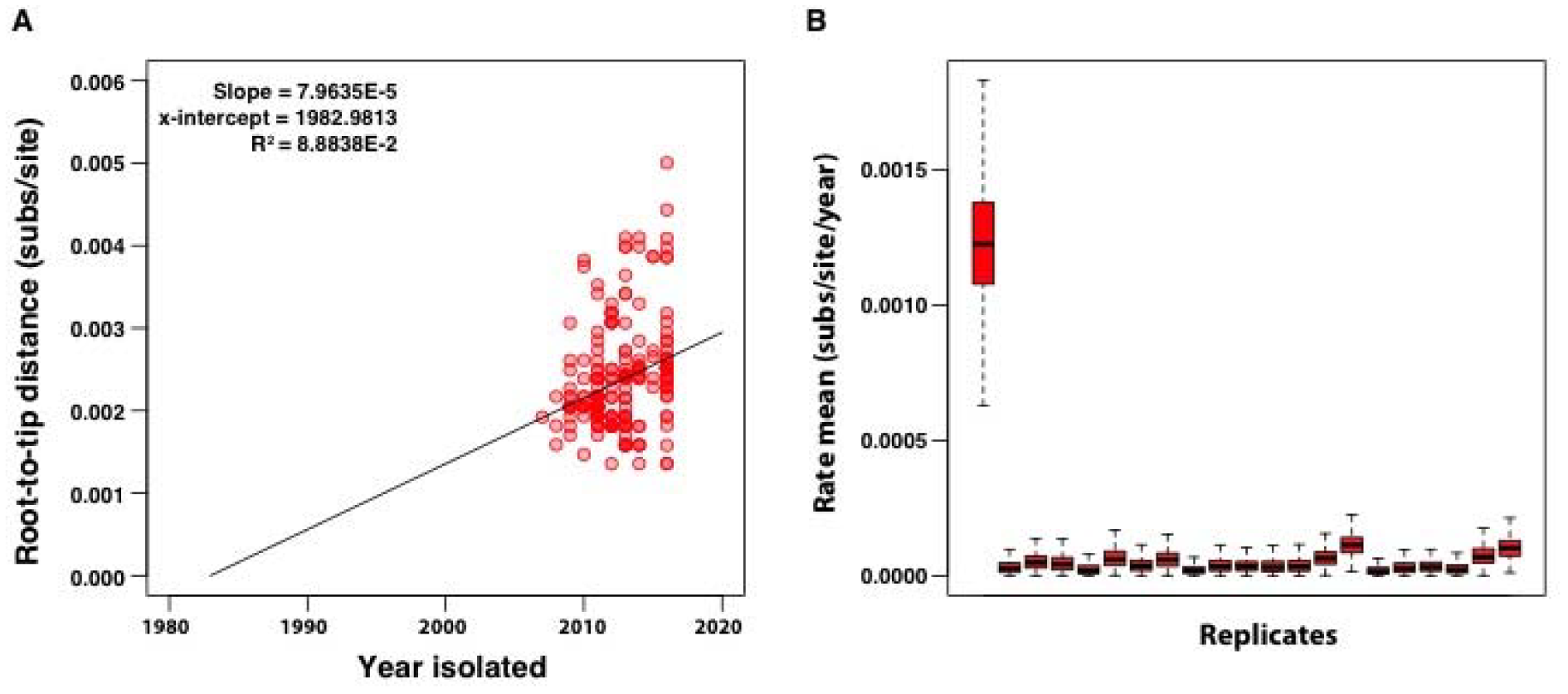
Temporal analysis of Nepalese H58 (4.3.1). (A) Tempest regression of root-to-tip distance as (in the SNP alignment) a function of sampling time, with the root of the tree selected using heuristic residual mean squared (each point represents a tip of the maximum likelihood tree). The slope is a crude estimate of the substitution rate for the SNP alignment, the x-intercept corresponds to the e of the root node, and the R^2^ is a measure of clocklike behaviour (B) Date randomisation test with the left most box plot showing the posterior substitution rate estimate from the SNP alignment of the data with the correct sampling times, and the remaining 20 boxplots showing the posterior distributions of the rate from replicate runs using randomised dates. The data are considered to have strong temporal structure if the estimate with the correct sampling times does not overlap with those from the randomisations.

**Supplementary Figure 3.**
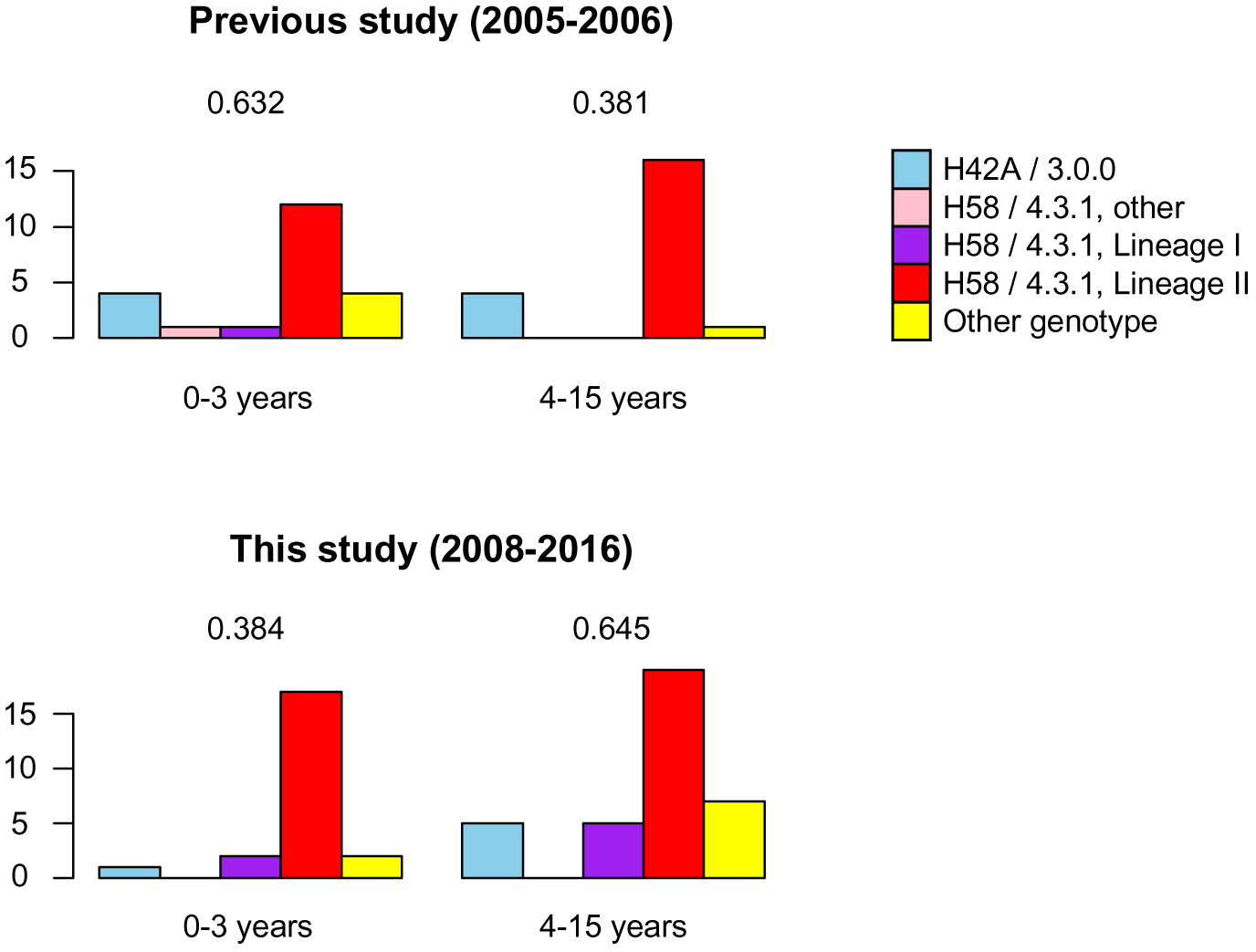
Direct comparison of *S*. Typhi genotype diversity detected in this study compared to an earlier study of children at the same hospital (Holt et al, 2010; ref 5), stratified by age group. Simpson’s diversity of *S*. Typhi genotypes is printed on each plot.

